# Piezo2 mechanosensitive ion channel is located to sensory neurons and non-neuronal cells in rat peripheral sensory pathway: implications in pain

**DOI:** 10.1101/2021.01.20.427483

**Authors:** Seung Min Shin, Francie Moehring, Brandon Itson-Zoske, Fan Fan, Cheryl L. Stucky, Quinn H. Hogan, Hongwei Yu

## Abstract

Piezo2 mechanotransduction channel is a crucial mediator of sensory neurons for sensing and transducing touch, vibration, and proprioception. We here characterized Piezo2 expression and cell specificity in rat peripheral sensory pathway using a validated Piezo2 antibody. Immunohistochemistry using this antibody revealed Piezo2 expression in pan primary sensory neurons (PSNs) of dorsal rood ganglia (DRG) in naïve rats, which was actively transported along afferent axons to both central presynaptic terminals innervating the spinal dorsal horn (DH) and peripheral afferent terminals in skin. Piezo2 immunoreactivity (IR) was also detected in the postsynaptic neurons of the DH and in the motor neurons of the ventral horn, but not in spinal GFAP- and Iba1-positive glia. Notably, Piezo2-IR was clearly identified in peripheral non-neuronal cells, including perineuronal glia, Schwann cells in the sciatic nerve and surrounding cutaneous afferent endings, as well as in skin epidermal Merkel cells and melanocytes. Immunoblots showed increased Piezo2 in DRG ipsilateral to plantar injection of complete Freund’s adjuvant (CFA), and immunostaining revealed increased Piezo2-IR intensity in the DH ipsilateral to CFA injection. This elevation of DH Piezo2-IR was also evident in various neuropathic pain models and monosodium iodoacetate (MIA) knee osteoarthritis (OA) pain model, compared to controls. We conclude that 1) the pan neuronal profile of Piezo2 expression suggests that Piezo2 may function extend beyond simply touch/proprioception mediated by large-sized low-threshold mechanosensitive PSNs, 2) Piezo2 may have functional roles involving sensory processing in spinal cord, Schwann cells, and skin melanocytes, and 3) aberrant Piezo2 expression may contribute pain pathogenesis.

## 1. Introduction

Mechanosensitive ion channels are a diverse population of ion channels that integrate various internal and external mechanical cues to electrochemical mechanotransduction signals with different biophysical properties and biological significance [16, 29, 39, 47]. The evolutionarily conserved Piezo channels, including Piezo1 and Piezo2, are mechanically activated ion channels that play a critical role in a variety of mechanotransduction processes of different cell types [15, 42, 76]. Piezo1 mediates shear stress and stretch-induced transmembrane currents mainly in nonneuronal cells, whereas Piezo2 have been found predominantly expressed in primary sensory neurons (PSNs), where it confers proprioception and touch perception, and as shown more recently, detection of noxious mechanical stimuli [43]. In humans, mutations of genes expressing these channels are not life-threatening but result in different pathologies [12, 21]. Lack of painful reactions to innocuous touch after skin inflammation is reported in the loss-of-function mutations of *PIEZO2* but not *PIEZO1* [12, 45, 49, 67].

Various techniques have been employed to define Piezo2 expression in the PSNs of DRGs and trigeminal ganglia (TG), all of which detect Piezo2 expression in the PSNs. Initially, *in situ* hybridization (ISH) revealed Piezo2 expression in a subpopulation (20∼60%) of large diameter low threshold mechanoreceptive (LTMR) PSNs in mouse DRG. However, recent studies also using ISH report that Piezo2 transcripts are expressed in all DRG-PSNs, including most notably the large diameter neurons implicated in mediating touch and proprioception, as well as small-sized nociceptive PSNs [4, 10, 15, 43, 48, 70, 71]. Similarly, single-cell RT-PCR reports that *Piezo2* mRNA is detected in 39% of all DRG-PSNs, while new efforts by single-cell RNA sequencing (RNAseq) identify Piezo2 in the majority of mouse DRG-PSNs [51, 72, 83]. Immunohistochemistry (IHC) has observed Piezo2 expression in up to 45∼80% of total PSNs of mouse DRG with high expression in large diameter PSNs, while a recent report describes Piezo2 expression in all PSNs of mouse DRG neurons, including large-diameter myelinated neurons encompassing LTMRs, as well as medium- and small-diameter nociceptors [43, 62, 69, 73]. These reports of Piezo2 expression in ganglia are derived from mouse or duck [56], but Piezo2 expression has not been systematically examined in the rat peripheral nervous system (PNS).

Piezo2-knockout mice fail to develop allodynia after skin inflammation [62], while upregulation of Piezo2 activity correlates with inflammation-induced pain states, and inflammatory signals are found to enhance Piezo2-mediated mechanosensitive currents [18]. PSN-specific Piezo2 knockout mice develop less sensitization to noxious mechanical stimulation following spared nerve injury [43], suggesting Piezo2 participation in detection of such stimuli in inflammatory and neuropathic pain settings [26]. In order to pursue the therapeutic potentials targeting Piezo2 for chronic pain in preclinical rat models, this study was designed to characterize Piezo2 expression in the PNS of rat. We found that Piezo2 was expressed by all DRG neurons, variety of non-neuronal cells in the peripheral sensory pathway, and spinal cord neurons. Presence of Piezo2 in spinal cord and peripheral glial cell populations suggests that Piezo2 may participate in sensory processing in spinal cord neurons and peripheral glial cells.

## 2. Methods

### 2.1. Animals

Adult male Sprague Dawley (SD) rats (6-8 week old, Charles River Laboratories, Wilmington, MA) were used. All animal experiments were performed with the approval of the Zablocki VA Medical Center Animal Studies Subcommittee and Medical College of Wisconsin Institutional Animal Care and Use Committee in accordance with the National Institutes of Health Guidelines for the Care and Use of Laboratory Animals. Animals were housed individually in a room maintained at constant temperature (22±0.5°C) and relative humidity (60±15%) with an alternating 12h light-dark cycle. Animals were given access to water and food *ad libitum* throughout the experiment, and all efforts were made to minimize suffering and the numbers of animal used. For tissue harvest euthanasia, animals were deeply anesthetized by isoflurane followed by decapitation with a well-maintained guillotine. The numbers of rats used are detailed in the relevant sections of the experiments.

### 2.2. Pain models

#### 2.2.1. Intraplantar CFA inflammatory pain

Inflammatory pain was induced by injection of a single dose of 100μl of CFA (Sigma-Aldrich, St. Louis, MO) at a concentration of 1.0 mg/ml or saline subcutaneously in the plantar surface of the right hind paw of isoflurane anesthetized rats, performed as described previously [24].

#### 2.2.2. Mechanical allodynia and hyperalgesia

Mechanical allodynia was assessed as the mechanical withdrawal threshold (von Frey, vF) and hyperalgesia was identified by noxious punctate mechanical stimulation (Pin test), as described previously [20]. In brief, vF test was performed by applying the calibrated monofilaments (Patterson Medical, Bolingbrook, IL) to the plantar surface of the hindpaw. Beginning with the 2.8 g filament, if a response was observed, the next smaller filament was applied, and if no response was observed, the next larger was applied, until a reversal occurred, defined as a withdrawal after a previous lack of withdrawal, or vice versa. Following a reversal event, four more stimulations were performed following the same pattern. The forces of the filaments before and after the reversal, and the four filaments applied following the reversal, were used to calculate the 50% withdrawal threshold [20]. Rats not responding to any filament were assigned a score of 25 g. Pin test was performed using the point of a 22 g spinal anesthesia needle that was applied to the center of the hindpaw with enough force to indent the skin but not puncture it. Five applications were separated by at least 10s each, which was repeated after 2 min, making a total of 10 touches. For each application, the induced behavior was either a very brisk, simple withdrawal with immediate return of the foot to the cage floor, or a sustained elevation with grooming that included licking and chewing, and possibly shaking, which lasted at least 1s. This latter behavior was referred to as hyperalgesic behavior [74], which is specifically associated with place avoidance. Hyperalgesia was quantified by tabulating hyperalgesia responses as a percentage of total touches.

#### 2.2.3. Tissue harvest for immunohistochemistry (IHC) and western blots

Six naïve rats with normal mechanical sensory thresholds by vF and Pin tests were used for IHC characterization of Piezo2 expression. Specifically, lumbar (L) 4 and 5 DRG, trigeminal ganglia (TG), lumbar segment spinal cord, sciatic nerve trunk segment proximal to the sciatic bifurcation, brain, and hindpaw glabrous and hair skin tissues, as well as lumbar spinal cord from CFA rats, were dissected and fixed in Richard-Allan Scientific™ Buffered Zinc Formalin (ThermoFisher, Rockford, IL) overnight (∼15 hr) for DRG, sciatic nerves, and skins; and 24 hr for spinal cord and brain; followed by processing for paraffin embedment. Serial sections at 5μm thickness; orientated at coronal for brain, sagittal for DRG and sciatic nerve, transverse for spinal cord, and perpendicular to the skin surface for glabrous and hair skin tissues; were prepared and mounted on the positive charged SuperForst microscope slides (ThermoFisher). For cryosection, fixed sciatic nerve tissues were cryoprotected by overnight infiltration with 30% sucrose solution in 1x phosphate-buffered saline (PBS), followed by mounting in OCT embedding compound, and freeze at −80 °C. Fifteen μm thickness cryosection were prepared on a crystate (Leica CM1950, Kyoto, Japan) and air-dried for 24 hr. For western blot experiments, L4/L5 DRG from CFA- and saline-injected rats, after cardiac perfusion with cold PBS, were dissected, snap frozen in liquid nitrogen, and stored at −80°C for extraction of protein.

### 2.3. Cell cultures

#### 2.3.1. *DRG dissociated culture and neuron-free SGC isolation* [78, 80]

In brief, the L4/5 DRG were rapidly harvested from the isoflurane anesthetized naïve rats and were incubated in 0.01% blendzyme 2 (Roche Diagnostics, Madison, WI) for 30 min, followed by incubation in 0.25% trypsin and 0.125% DNase for 30 min; both dissolved in Dulbecco’s modified Eagle’s medium/F12 (DMEM/F12) with glutaMAX (ThermoFisher). After exposure to 0.1% trypsin inhibitor and centrifugation, the pellet was gently triturated and dissociated cells cultured in Neural basal media A (ThermoFisher) plus 0.5 μM glutamine at 37°C in humidified 95% air and 5% CO_2_. Neuron-free SGC culture was established by a differential attachment protocol for SGC isolation, as we described previously [58].

#### 2.3.2. Sciatic nerve (SN) Schwann cell isolation

the relevant steps required for nerve processing, enzymatic dissociation, and cell plating using the SN from one adult male rat was previously described, with minor modifications [5]. In brief, bilateral sciatic nerves were harvested and attached adipose and muscular tissue stripped off using fine forceps. Subsequently, the outermost epineurial layer and the epineurium were removed as one single sheath and collected for enzymatic digestion. The fibers were extensively teased until no individual fascicles were evident. The epineurium and teased fibers were subjected to digestion with an enzymatic cocktail for 4hr as described in DRG dissociated culture. The end products of enzymatic digestion were filtered and subsequently collected by centrifugation.

#### 2.3.3. Cell lines

As a second source of Schwann cells, primary Schwann cells isolated from human spinal nerve were obtained from Neuromics (HMP303, Edina, MN). Primary adult human epidermal melanocytes were purchased from ThermoFisher (C0245C). Human SCs and isolated rat SCs were cultured in Schwann cell medium (Edina), and human epidermal melanocytes cultured in Medium 254 with melanocyte growth supplement (ThermoFisher), according manufacturer’s protocols. Neuronal cell lines of Neuro2A (N2A, mouse brain cortex neurons), F11 (rat DRG neuron-like cells), and Neuroblastoma B104 cells (B104) were obtained from ATCC (Manassas, VA), and rat DRG-neuron 50B11 cells (50B11) were as prior reported [78], and these cells were cultured by a standard protocol at 37 °C with 5 % CO_2_ in DMEM supplemented with 10% FBS and 1% penicillin-streptomycin (ThermoFisher).

### 2.4. *In vitro* Piezo2 knockdown for Piezo2 antibody validation

For validation of Piezo2 antibody in a genetic strategy, the plasmids expressing of a short hairpin RNA against mouse Piezo2 in the coding region and a scrambled control (SC), in silico designed by use of BLOCK-iT RNAi Designer (Invitrogen), were generated. Specifically, a designed convergent U6 and H1 promoters driven apposing piezo2-shRNA (or SC) expression cassette was synthesized (Genscript, Piscataway, NJ) and cloned into Mlul site of an expression plasmid in which EGFP was transcribed by a CMV promoter. These plasmids were named as pCMV-Piezo2shRNA-CMV-EGFP (pCMV-Piezo2shRNA) and pCMV-Piezo2SC-CMV-EGFP (pCMV-Piezo2SC), respectively. Transfection on N2A cells was performed by a standard PEI transfection protocol and Piezo2 knockdown efficacy determined by qPCR, western blot, and Piezo2 channel patch recording was performed 48hr post-transfection.

### 2.5. RT-PCR and quantitative PCR (qPCR)

Total RNA was extracted from culture cells and DRG using RNAeasy kit (Qiagen, Carlsbad, CA, USA) and then treated with DNase I (Life Technologies). The concentration and purity of the total RNA were evaluated with a spectrophotometer. Complementary DNA (cDNA) was synthesized from 1.0 μg RNA using the Superscript III first strand synthesis kit with random hexamer primers (Life Technologies). PCR was carried out to determine Piezo2 mRNA in 50B11 cells, DRG, and isolated SGCs and Schwann cells on a Bio-Rad C1000 PCR machine and specific intron-spanning primers from Piezo2 (and Piezo1) (**Table 1**). All of the primers were designed by use of MacVector version17.0.8 (https://www.macvector.com) and synthesized by Integrated DNA Technologies, Inc. (http://www.idtdna.com). The thermal cycling conditions were one cycle at 95°C for 3 min, 40 cycles at 95°C for 10 s, 60°C for 30 s, and 72°C for 50 s, followed by one cycle at 72°C for 5 min. A negative control with double-distilled water was performed for each PCR (and qPCR) experiment. QPCR was performed to determine Piezo2 knockdown effects by transfection of piezo2-shRNA into N2A cells using IQ Syber Green supermix (Bio-rad, Hercules, CA, USA) on a Bio-rad CFX96 Real-time PCR Machine and specific intron-spanning primers to quantify the cDNA levels of Piezo2 (**Table 1**). The thermal cycling conditions were one cycle at 95°C for 2 min, 40 cycles at 95°C for 10 s, 60°C for 10 s. The efficiency of the primers was 92–100% with r2>0.990. For each sample, two inter-run determinations were carried out, and two replicates in each run were averaged. For the comparative CT method, GAPDH were used as the housekeeping gene for calculation of fold differences in expression of Piezo2 mRNA. The relative amount of mRNA transcript in N2A cells transfected by Piezo2-shRNA to scrambled controls was measured by 2^− ΔΔC^ t calculation, where ΔΔC_t_ = (C_t piezo2_ − C_t Gaphd_)_contorl_ − (C_t piezo2_ − C_t Gapdh_) _Piezo2shRNA_

**Table 1.**
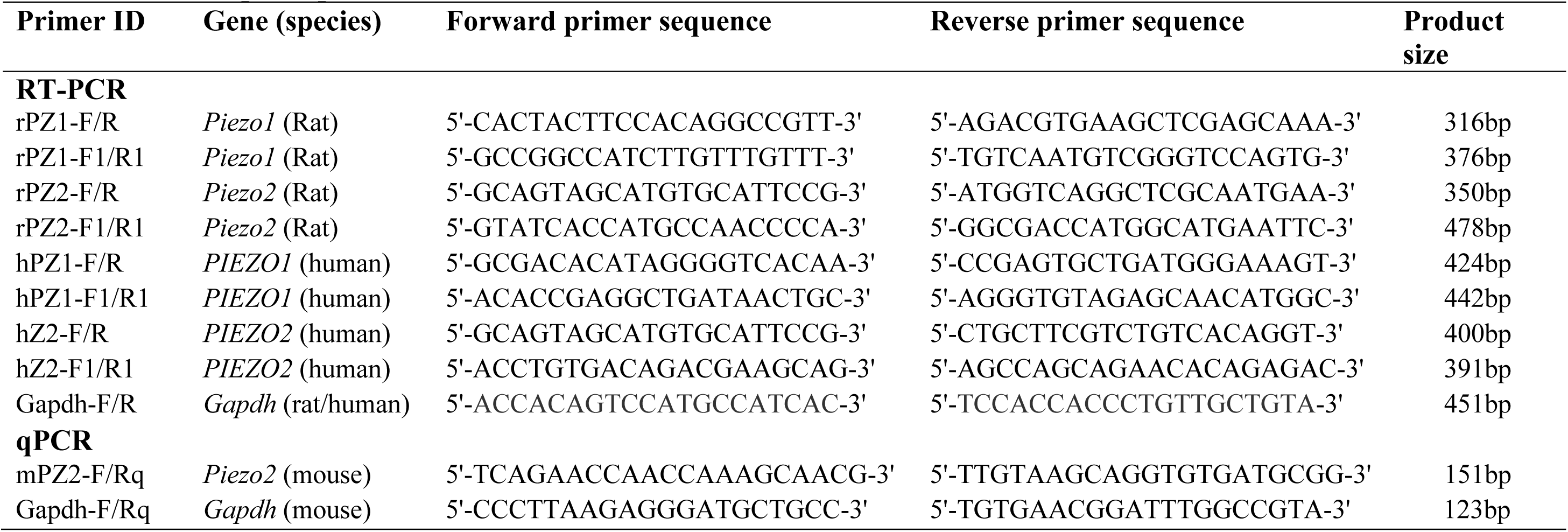
PCR and qPCR primers

### 2.6. Patch clamp recordings

For validation of Piezo2 functional knockdown, N2A cells transfected by pCMV-Piezo2shRNA or scrambled RNA (scrambled) were used for recording mechanically activated currents under the conventional whole-cell configuration. N2A cells were superfused continuously with RT extracellular normal HEPES solution containing (in mM): 140 NaCl, 5 KCl, 2 CaCl2,1 MgCl2, 10 HEPES, and 10 glucose, pH 7.4 ± 0.05, and 310 ± 3 mOsm, and viewed on a Nikon Eclipse TE200 inverted microscope. Cells were patch clamped in voltage clamp with a borosilicate glass pipette (Sutter Instrument Company, Novato, CA) filled with intracellular normal HEPES solution containing (in mM): 135 KCl, 10 NaCl, 1 MgCl2, 1 EGTA, 0.2 NaGTP, 2.5 ATPNa2, and 10 HEPES, pH 7.20 ± 0.05, and 290 ± 3 mOsm. Cell capacitance and series resistance were maintained below 10 MΩ. Mechanical stimulation was elicited using a second borosilicate glass pipette that was driven by a piezo stack actuator (PA25, PiezoSystem Jena, Jena, Germany) at a speed of 106.25 μm/ms. Cells were stimulated with increasing displacements of 1.7 μm/Volt for 200 ms, and 2 min rest was allowed between displacements to avoid sensitization/desensitization of the cell membrane. Data was recorded using PatchMaster via an EPC10 amplifier HEKA Electronics, Holliston, MA). Data were analyzed using FitMaster (HEKA Electronics).

### 2.7. Immunofluorescent staining

#### 2.7.1. Immunocytochemistry (ICC) and immunohistochemistry (IHC)

ICC on cultured human and isolated rat Schwann cells, human melanocytes, and DRG dissociated cultures, as well as IHC on tissue sections, were performed according to standard procedures [78]. For ICC, the cultured cells were fixed in 2% paraformaldehyde (PFA) in 1xPBS solution for 5 min; and for IHC, the formalin-fixed, paraffin-embedded (FFPE) tissue sections were deparaffinized, hydrated, and treated by heat-induced antigen epitope retrieval in 10mM citrate buffer, pH 6.0. Cryosections were treated by 3% hydrogen peroxide for 10 min to block endogenous peroxidase activity. Non-specific binding was reduced by incubating the sections for 30 min with a solution of 5% BSA in PBS plus 0.05% Tween20 (PBST) solution. Samples were first immunolabeled with the selected primary antibodies in a humid atmosphere overnight at 4°C (**Table 2**). All antibodies were diluted in PBST, containing 0.05% Triton X-100 and 5% bovine serum albumin (BSA). Normal immunoglobulin G (IgG from the same species as the first antibody, **Table 2**) was replaced for the first antibody as the negative controls. The appropriate fluorophore-conjugated (Alexa 488 or Alexa 594, 1:2000) secondary antibodies (Jackson ImmunoResearch, West Grove, PA) were used to reveal immune complexes. Afterward, the sections were rinsed for 10 min in PBS and either processed for a colabeling of primary and secondary antibodies or coverslipped under Shur/Mount mounting medium (ThermoFisher). To control for false-positive results attributable to cross-binding in double-label combinations, each primary antibody raised in a different species was used. To stain nuclei, 1.0μg/ml Hoechst33342 (Hoechst, ThermoFisher) was added to the secondary antibody mixture. The immunostaining was examined, and images captured using a Nikon TE2000-S fluorescence microscope (El Segundo, CA) with filters suitable for selectively detecting the green and red fluorescence using a QuantiFire digital camera (Optronics, Ontario, NY). For double label colocalization, images from the same specimen but showing different antigen signals were overlaid by digitally merging the captured images.

**Table 2.** Primary antibodies and IgG controls used in this study

| **Antibody<sup>a</sup>** | **Host** | **Supplier/Cat#<sup>b</sup>** | **Dilution** |
| --- | --- | --- | --- |
| IB4 |  | LF/I21413 | 1.0µg/ml |
| Piezo2 | Rabbit polyclonal | Prosci/8613 | 1:100 (IHC), 1:1000 (Wb) |
| Piezo2 | Rabbit polyclonal | Alomone/APC090 | 1:100 (IHC), 1:1000 (Wb) |
| CGRP | Mouse monoclonal | SCB/sc57053 | 1:600 (IHC) |
| GFAP | Rabbit polyclonal | Dako/Z0334 | 1:1000 (IHC) |
| GFAP(GA5) | Mouse monoclonal | CS/3655 | 1:200 (IHC) |
| Tubb3 | Mouse monoclonal | SCB/sc80016 | 1:500 (IHC), 1:1000 (Wb) |
| NKA1α | Mouse monoclonal | SCB/sc514614 | 1:1000 (IHC) |
| TrkB | Mouse monoclonal | SCB/sc136990 | 1:500 (IHC) |
| GS | Mouse monoclonal | SCB/sc74430 | 1:800 (IHC) |
| S100 | Mouse monoclonal | TF/MA5-12969 | 1:500 (IHC) |
| NF200 | Mouse monoclonal | Sigma/N5389 | 1:1000 (IHC) |
| Syp | Mouse monoclonal | SCB/sc17750 | 1:200 (IHC) |
| Synpr | Mouse monoclonal | SCB/sc376761 | 1:200 (IHC) |
| CK14 | Mouse monoclonal | SCB/sc53253 | 1:200 (IHC) |
| MBP | Mouse monoclonal | CS/41168 | 1:800 (IHC) |
| P75NTR | Mouse monoclonal | SCB/sc271708 | 1:100 (IHC) |
| Iba1 | Rabbit polyclonal | Wako/019-19741 | 1:1000 (IHC) |
| Iba1 | Mouse monoclonal | Sigma/ MABN92 | 1:200 (IHC) |
| GAPDH | Mouse monoclonal | Sigma/SAB1403850 | 1:5000 (Wb) |
| IgG control | Mouse | LF/31903 | 1:100~400 |
| IgG control | Rabbit | LF/MA5-16384 | 1:100~1000 |

<sup>a</sup> Antibody abbreviations IB4, Isolectin IB4; Piezo2, Piezo type mechanosensitive ion channel component 2; CGRP, calcitonin gene-related peptide; GFAP, Glial fibrillary acidic protein; Tubb3, β3-Tubulin; NKA1α, sodium/potassium ATPase 1α; TrkB, tropomyosin receptor kinase B; GS, glutamine synthetase; NF200, neurofilament-200; Syp, synaptophysin; Synpr, synaptoporin; CK14, cytokeratin 14; MBP, myelin basic protein; P75NTR, neurotrophin receptor p75; GAPDH, glycolytic enzyme glyceraldehyde-3-phosphate dehydrogenase
<sup>b</sup> LF, Life Technologies, Carlsbad, CA; Alomone, Alomone Labs, Jerusalem, Israel; SCB, Santa Cruz Biotechnology, Santa Cruz, CA; CS: Cell signaling, Danvers, MA; Wako, Richmond, VA; Dako: Carpinteria, California; Sigma, Sigma-Aldrich, St Louis, MO

Two independently developed polyclonal Piezo2 antibodies, purchased from ProSci (Poway, CA) and Alomone (Jerusalem, Israel), were used. For control purposes (Piezo2 immunostaining), representative sections were processed in the same way as described but using nonimmune rabbit sera instead of the Piezo2 primary antibody in the incubation. We also tested the specificity by preincubating the antibody solution with the manufacturer’s specific PIEZO2 antigenic peptides (2 μg/ml) for 2 hr prior to immunostaining, as described previously [68]. The specificities of the other antibodies used in this study have been previously confirmed, and the specificity of secondary antibodies was tested with omission of the primary antibodies, which always resulted in no immunostaining [68, 75, 78, 79].

#### 2.7.2. Measurement and quantification of immunostaining

Positive marker antibody immunostainings were defined as the cells with the fluorescence intensity greater than average background fluorescence plus 2 standard deviations of the cells in an adjacent section in the same slide of negative control (first antibody omitted) under identical acquisition parameters (n=10 for different markers), identified by Hoechst counterstain at a different wavelength [77].

Intensity correlation analysis (ICA) was performed to determine colocalization of Piezo2 with neuronal plasma membrane (PM) marker sodium/potassium ATPase 1 alpha (NKA1a) and SGC marker glial fibrillary acidic protein (GFAP), as previously described using an ImageJ 1.46r software plugin colocalization analysis module (http://imagej.nih.gov/ij) [35, 58, 80]. In brief, fluorescence intensity was quantified in matched region of interests (the green and red colors varied in close synchrony) for each pair of images. Mean background was determined from areas outside the section regions and was subtracted from each file. On the basis of the algorithm, in an image where the intensities vary together, the product of the differences from the mean (PDM) will be positive. If the pixel intensities vary asynchronously (the channels are segregated), then most of the PDM will be negative. The intensity correlation quotient (ICQ) is based on the nonparametric sign-test analysis of the PDM values and is equal to the ratio of the number of positive PDM values to the total number of pixel values. The ICQ values are distributed between **-**0.5 and **+**0.5 by subtracting 0.5 from this ratio. In random staining, the ICQ approximates 0. In segregated staining, ICQ is less than 0, while for dependent staining, ICQ is greater than 0.

Lumbar spinal cord sections from control rats and from rats subjected to CFA inflammatory pain, as well as archival spinal cord sections from neuropathic pain models of SNL, SNI, and TNI (4wk post injury), and MIA-OA pain (5wk after MIA knee injection), as previously reported [37, 57-59, 80, 81], were used to evaluate whether pain is associated with alteration of Piezo2 expression in the DH. For quantification of DH piezo2 immunostaining, the Image J v.1.46 was used to quantify changes in immunolabeled fluorescent intensities as described previously [14, 57], with some minor modifications. In brief, the sections with symmetrical width of DH throughout the mediolateral axis were used for measurement, and the upper and lower threshold optical intensities were adjusted to encompass and match the immunoreactivity (IR) that appears in red. A standardized rectangle was first positioned over laminae territory throughout the mediolateral axis on the contralateral DH. The area and intensity of pixels within the threshold value representing IR were calculated and the integrated intensity was the product of the area and density. The same box was then moved to the corresponding position on the opposite DH and the integrated intensity of pixels within the same threshold was again calculated. Comparisons of both sides of DHs were made only within the same sections and intensity values on the ipsilateral side were expressed as a percent of the contralateral side, providing an estimate of fold change (ratio of ipsilateral/contralateral) for each section.

### 2.8. Immunoblot analysis of Piezo2 expression

Cell and tissue lysates were prepared from Neuro2A, F11, B104, 50B11 cells, and DRG. The collected cell pellets and pooled L4/L5 DRG from CFA and control rats were extracted using 1x RIPA ice-cold buffer (20 mm Tris-HCl pH 7.4, 150 mm NaCl, 1% Nonidet P-40, 1% sodium deoxycholate, 0.1% SDS, with 0.1% Triton X100 and protease inhibitor cocktail) and rotated at 4°C for 1 h before the supernatant was extracted by centrifugation at 12,000 g at 4°C for 5 min. To examine the subcellular localization of Piezo2, DRG tissues were homogenized and then fractionated to obtain plasma membrane and cytosolic fractions, using the ProteoExtract Subcellular Proteome Extraction Kit (Millipore, Billerica, MA), according to the manufacturer’s instructions. This kit contains extraction buffers with ultrapure chemicals to ensure high reproducibility, protease inhibitor cocktail to prevent protein degradation, and benzonase nuclease to remove contaminating nucleic acids. Protein concentration was determined using Pierce BCA kit (ThermoFisher). Equivalent protein samples (cell lysates and DRG fractionized membrane and cytosol) were size separated using 10% or 4-20% SDS-PAGE gels (Bio-Rad), transferred to Immun-Blot PVDF membranes (Bio-Rad), and blocked for 1 hr in 5% skim milk. In some experiments, the transferred PVDF membranes were cut into two halves along protein size around 70KDa and were subsequently incubated overnight at 4°C with appropriate antibodies. Immunoreactive proteins were detected by Pierce enhanced chemiluminescence (ThermoFisher) on a ChemiDoc Imaging system (Bio-Rad) after incubation for 1 hr with HRP-conjugated second antibodies (1:5000, Bio-Rad). Between each step, the immunoblots were rinsed with Tris-buffered saline containing 0.02% Tween-20 (TBST). To verify the band specificity of Piezo2 detection using a rabbit Piezo2 antibody, the antibody solution was preincubated with the manufacture’s specific Piezo2 antigenic peptides (2 μg/ml) for 2 hr prior to immunoblotting.

### 2.9. Statistics

Statistical analysis was performed with GraphPad PRISM 8 (GraphPad Software, San Diego, CA). Significances of ICQs of Piezo2 immunocolocalization with NKA1*α* and GFAP were analyzed by means of the normal approximation of the nonparametric Wilcoxon rank test, as described previously [35, 80]. Mechanical allodynia and hyperalgesia were compared between groups by Student’s *t* test for von Frey and by Mann–Whitney test for Pin. Patch recordings were analyzed using a repeated measures two-way ANOVA with Sidak *post-hoc* comparison. The differences of the targeted gene expression by qPCR, immunoblots, and DH Piezo2-IR intensity analysis on the spinal cord sections between different groups were compared with 2-tailed unpaired Student’s t test or Mann–Whitney test where appropriate. Results are reported as mean and SEM. P<0.05 were considered statistically significant.

## 3. Results

### 3.1. Specificity determination of Piezo2 antibody

Since the specificity and sensitivity for detection of Piezo2 expression is largely dependent on the quality of the first antibody, we decided to use two different, independently developed Piezo2 antibodies to optimize immunological detection of Piezo2 for ensuring specific detection of the Piezo2 protein expression in the cultured cells and DRG tissue. These two Piezo2 antibodies were chosen because the manufacturer’s western blot data show detection of protein bands with molecular mass of putative canonical Piezo2 and its isoforms (GenBank and ExPASy database), as well as the antigenic peptides are available. The first Piezo2 antibody was purchased from ProSci (Fort Collins, CO), which is an affinity chromatography purified rabbit polyclonal antibody raised against a intracellular peptide corresponding to 19 amino acids (HLTASLEKPEVRKLAEPGE, 14/19 homologous to rat sequence) near the amino terminus of human PIEZO2 (NP_071351), and predicted detection of PIEZO2 proteins of 233 and 305 KDa by immunoblot, no cross-reaction to Piezo1, and signals are blocked by antigenic peptide preabsorption. The second Piezo2 antibody, obtained from Alomone (Jerusalem, Israel), is also a rabbit polyclonal antibody which is raised, and affinity purified on immobilized antigenic peptide (RTIFHDITRLHLD, 12/13 homologous to rat sequence) corresponding to 13 amino acid residues (1092 -1104) of human PIEZO2. Manufacturer’s in-house data show that this antibody detects Piezo2 expression in rat DRG lysates of ∼300 and ∼180 kDa by immunoblot and signals are blocked by antigenic peptide preabsorption. The antigenic peptide sequences of these two antibodies are located in the predicted intracellular loop regions of human Piezo2 [71].

An initial experiment was performed to characterize Piezo2 detection in the lysates prepared from several neuronal cell lines and from DRG of naïve rat by use of Piezo2 antibody from ProSci. The canonical Piezo2 is composed of ∼2800 amino acids with a molecular mass at ∼310KDa and predicted to be modified by phosphorylation and glycosylation. Also, PSN *Piezo2* has been found to undergoes differential gene splicing, resulting in many alternatively spliced and different functional Piezo2 mRNAs that may be translated to different molecular weight (MW) protein isoforms [7, 63]. Seven or eight potential Piezo2 protein isoforms are computationally mapped in human and mouse (https://www.uniprot.org); but whether they are the biological products derived from gene alternative splicing or canonical Piezo2 proteolytic processing are remained to be established. Multiple bands in the range of 310-150KDa were noted upon immunoblotting in the homogenates of neuronal cell lines, all of which were eliminated by preincubation with excess (2μg) manufacturer’s immunogenic peptide (**Fig. 1A**). In DRG lysate, the antibody detected several bands with predicted molecular masses approximately at 310, 240, 180, and 130 KDa, which we termed as putative “isoforms” 1-4 of the target protein, since they were all eliminated by preincubation with immunogenic peptide (**Fig. 1B**). To verify membrane localization, we performed Piezo2 immunoblots in the cytosolic and membrane fractions prepared from the DRG harvested from naïve rats and results showed that the putative Piezo2 isoform 2 (∼240KDa) and 3 (∼180KDa) were more enriched in membranes while the canonical Piezo2 isoform (KDa ∼310) and putative isoform 4 (KDa ∼130) were more enriched in cytosolic fractions (**Fig. 1C**).

**Figure 1.**
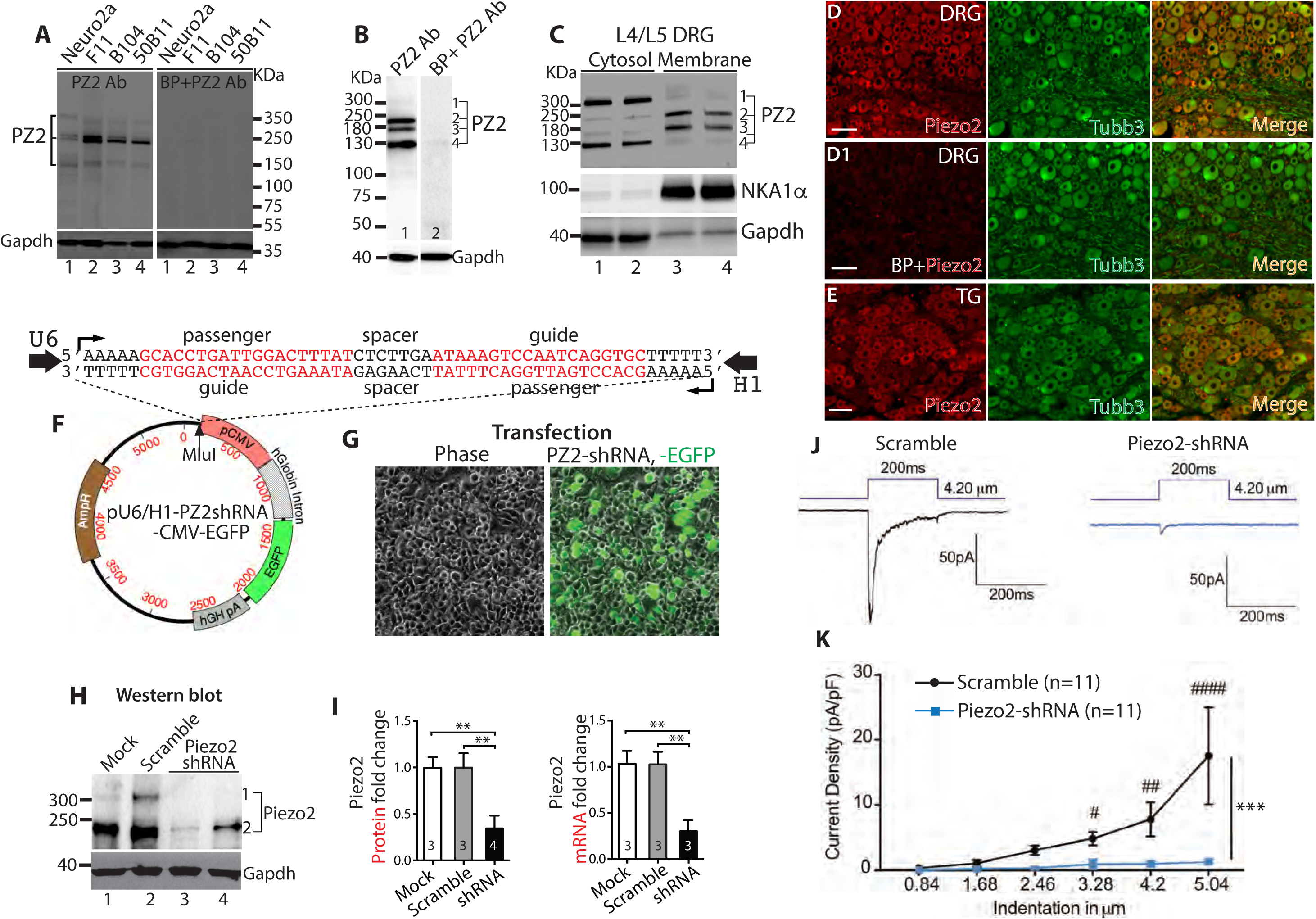
Specificity of Piezo2 (PZ2) antibody in detection of Piezo2 expression. Several bands around 310-150KDa are detected by immunoblot (IB) in the lysates of neuronal cell lines, which are eliminated by preincubation with blocking peptide (BP) (**A**). In DRG lysate, the antibody detects several bands at approximately 310, 240, 180, and 130 KDa and marked as putative “isoforms (iso)” 1-4, respectively, of the target protein since they are eliminated by preincubation with BP (**B**). Piezo2 IB in the cytosolic and membrane fractions prepared from pooled L4/L5 DRG show that the putative Piezo2 iso2 (∼240KDa) and iso3 (∼180KDa) are more enriched in membranes while the canonical Piezo2 (KDa ∼310) and putative iso4 (KDa ∼130) are more localized in cytosol (**C**). Representative images of double immunostaining (IS) of Piezo2 (red) and Tubb3 (green) on DRG sections reveal pan-neuronal Piezo2-IR (**D**) and preincubation with BP eliminated the staining (**D1**). Representative images of double-IS of Piezo2 (red) and Tubb3 (green) on trigeminal ganglia (TG) sections reveal pan-neuronal Piezo2-IR (**E**). Scale bars: 100 μm for all. Expression plasmid map (**F**) in which U6 and H1 promoters transcribe convergent apposing Piezo2-shRNA expression. The Piezo2-shRNA cassette (sequences shown on the top of plasmid map) is cloned into Mlul site (pointed by arrowhead) of plasmid and named pU6/H1-Piezo2shRNA-CMV-EGFP (Piezo2-shRNA). Transfection of Piezo2-shRNA into N2A cells yielded 70-80% transfection rate (n=4) (**G**); Piezo2 protein in N2A lysates analyzed by IB using ProSci Piezo2 antibody: Lane 1, mock transfection; lane 2, transfection with scrambled RNA; lane 3-4, duplicate transfection using Piezo2-shRNA (**H**); Densitometry of IBs of fold change in Piezo2 protein (left) and qPCR (right) show fold change in Piezo2 mRNA (**p<0.01, one-way ANOVA, Turkey *post-hoc* comparison) (**I**). Representative traces (**J**) and mean current density in response to mechanical force ramp (**K**) shows that Piezo2 knockdown significantly decreased mechanically-evoked rapid-adapt inward currents, compared to controls. ***p<0001; two-way ANOVA of main effects of groups with Bonferroni *post-hoc* comparison. ^#, ##^, and ^###^ denote p<0.05, <0.01, and<0.001, respectively, between groups.

IHC revealed a pan-neuronal pattern of Piezo2-IR in DRG and preincubation with Piezo2-specific antigenic peptide eliminated IHC staining (**Fig.1D, D1**). This effect was not observed when IHC was performed by preincubating Piezo2 antibody with a non-specific (Ca_V_3.2) antigenic peptide (Alomone; not shown), which provides additional validation for the selectivity. A pan-neuronal pattern of Piezo2-IR was also identified in the TG (**Fig. 1E**). Similar findings were confirmed using the second independent Piezo2 antibody (Alomone) by immunoblot and IHC on DRG tissues (see below), but this antibody seemed somewhat less sensitive for detecting Piezo2 by IHC. IHC using Alomone Piezo2 antibody on sections from TG sections also showed a pan-neuronal pattern of Piezo2 expression (**Suppl. Fig. 3F)**.

Selectivity of the ProSci Piezo2 antibody was further validated by comparing the relevant immunoblot Piezo2 protein level in the control N2A cells versus the N2A cells in which the RNAi induced Piezo2 knockdown, as was verified by qPCR and patch-recording of Piezo2 channel activity. As shown in **Fig. 1F-I**, shRNA-mediated knockdown of Piezo2 reduced the Piezo2-antibody recognized bands (∼310KDa and ∼250KDa), compared to the bands in the scrambled control-transfected cells, and these results correlate with an equivalent reduction of Piezo2 mRNA by qPCR quantification (approximately 80% for both; ∼310KDa and ∼250KDa bands in western blots were combined for calculation). In parallel, shRNA knockdown reduced mechanically-evoked rapid-adapting whole-cell current (**Fig. 1J, K**). These data provide evidence supporting the specificity of ProSci Piezo2 antibody. Based on these findings, the ProSci Piezo2 antibody was used for further IHC and immunoblot experiments, unless stated otherwise.

### 3.2. Piezo2 expression in the PNS

#### 3.2.1. Neuronal and glial Piezo2 expression in DRG (Fig. 2)

We next determined the Piezo2 expression within DRG sections by double immunolabeling using Piezo2 antibody with various established and distinct neuronal markers including Tubb3 (pan neurons), nonpeptidergic IB4-biotin (nonpeptidergic small neurons), peptidergic CGRP (peptidergic neurons), NKA1*α* (neuronal plasma membrane), NF200 for large A*β* low-threshold mechanoreceptors (A*β*-LTMRs) and proprioceptors, and TrkB (A*δ*-LTMRs) [34, 53, 83]. We found Piezo2-IR in all Tubb3-positive PSNs, suggesting pan-neuronal expression of Piezo2 in rat DRG, a finding that is consistent with the recent results from RNAscope ISH and IHC [69, 83]. All IB4- and CGRP-positive neurons which consist of 45% and 59% of Piezo2-positive neurons, respectively, were Piezo2 immunopositive, indicating that Piezo2 is highly expressed by unmyelinated (C-type) and thinly myelinated (Aδ-type) PSNs that convey the thermal and mechanoreceptive nociceptive signals generated at peripheral nerve terminals to neurons in lamina I-II of the spinal cord [17]. Nociceptive neurons in adult rat DRG can be double positive for both IB4 and CGRP (30-40%) [80], suggesting that there is a substantial proportion of small Piezo2-IR neurons positive for both IB4/CGRP. Colabeling of Piezo2 with NKA1*α* revealed enriched Piezo2 profiles in the PSN plasma membrane, especially those of larger diameter neurons. Nearly all NF200-neurons that consist of 30% among Piezo2-IR neurons were Piezo2 immunopositive, suggesting detection of Piezo2 expression in the A*β*-LTMRs and proprioceptive PSNs that transmit mechanoreceptive and proprioceptive signals via thickly myelinated afferents (Aβ-type) to spinal lamina III-V. Most of Piezo2-positive PSNs were also positive for TrkB, suggesting high expression of Piezo2 in the A*δ*-LTMR neurons (**Fig. 2A-F1**). To obtain further insight into plasma membrane (PM) localization of Piezo2, we examined the ICA of the images co-stained Piezo2 with NKA1*α*. Overlaid images of Piezo2 with NKA1*α* showed colocalizations of the two patterns of immunopositivity (**Fig. 2F1**), but the ICA plots of Piezo2 and NKA1*α* resulted, however, in a more complex relationship in which the data tended to cluster along both positive and negative axes with ICQ 0.04-0.2 (*p* <0.01∼ 0.001, n=10), indicating partial localization of Piezo2 in neuronal PM (**G**). ICC on DRG dissociated culture also identified Piezo2 expression in the neurons, as well as S100- and GFAP-positive glial cells (**Suppl. Fig. 1**).

**Figure 2.**
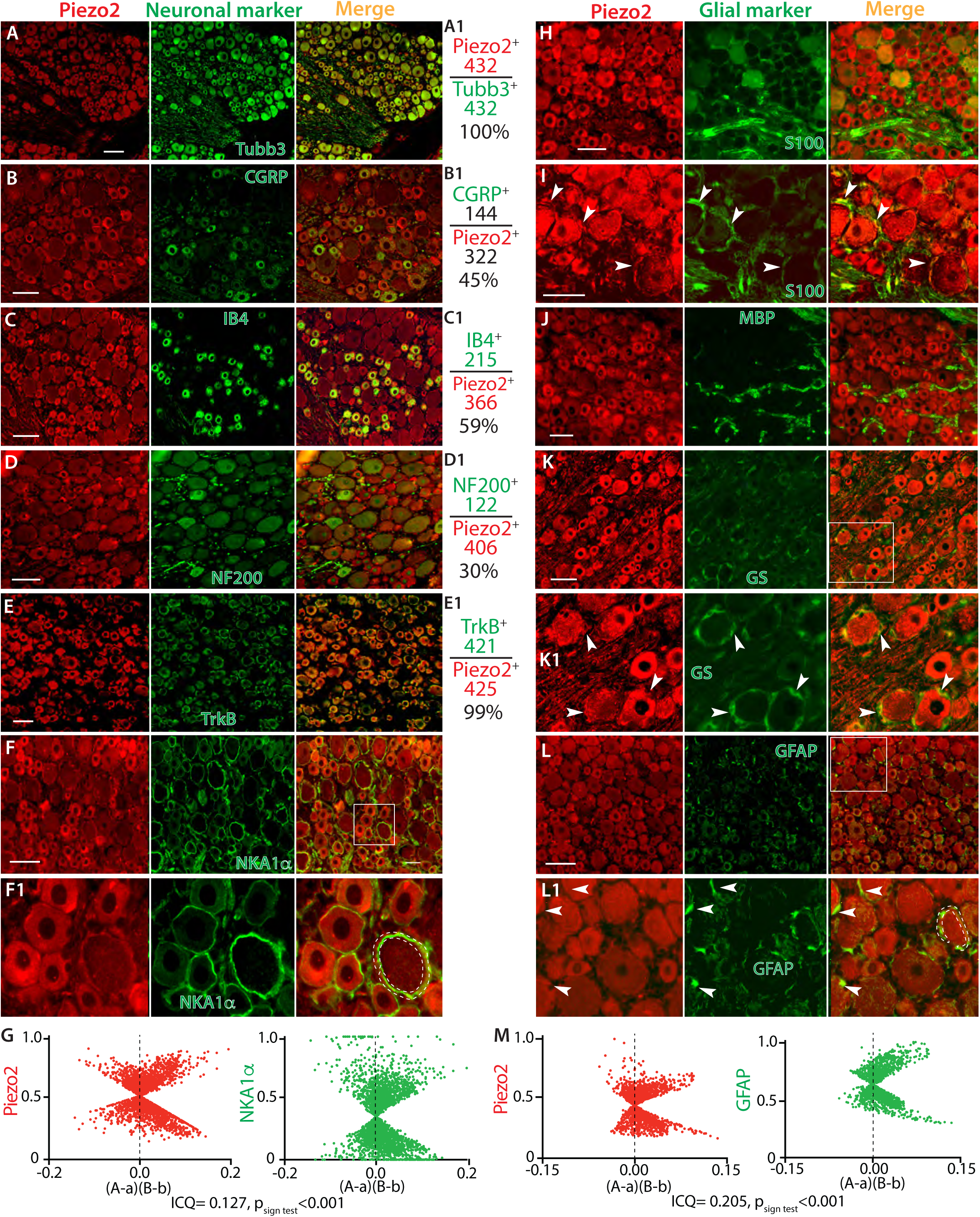
Double immunostaining (IS) of Piezo2 (PZ2) with a selection of neuronal and glial markers. Representative montage images of DRG sections show Piezo2-IR (red) co-stained with a selection of DRG neuronal markers (green), including Tubb3 (**A**), IB4 (**B**), CGRP (**C**), NF200 (**D**), TrkB (**E**), and NKA1*α* (**F**) with the squared region shown at high magnification (**F1**). The panels in the right-side of **A**-**E** calculate the percentage of Piezo2-neurons overlaid to Tubb3-neurons (**A**1), as well as neurons positive for IB4 (**B1**), CGRP (**C1**), NF200 (**D1**), and TrkB (**E1**) overlaid to Piezo2-neurons. The numbers are the counted Piezo2-IR neurons (red) and marker-labeled neurons (green). ICA analyzes colocalization between Piezo2 and NKA1*α* by an ImageJ 1.46r software plugin colocalization analysis module (**G**). Scatter plots for the region demarcated by the white dashed line in **F1** panel show data clustered along both positive and negative axes for both Piezo2 and NKA1*α*. “A” is the intensity of Piezo2 while “a” is the average of these values, and “B” is the intensity of NKA1a while “b” is the average of these values. For this region, the ICQ value is 0.127 (P_sign test_<0.001), indicating partial immunocolocalization. Representative montage images show Piezo2 (red) with a selection of glial cell markers (green), including S100 (**H, I**), MBP (**J**), GS (**K**) with the squared region shown at high magnification (**K1**), and GFAP (**L**) with the squared region shown at high magnification (**L1**). White arrowheads in panel **I, K1**, and **L1** point to the immune-colabeled glial cells. ICA analysis for colocalization between Piezo2 and GFAP for the region demarcated by the white dashed line in **L1** panel show scattered plot data clustered along both positive and negative axes for both Piezo2 and GFAP. “A” is the intensity of Piezo2 while “a” is the average of these values, and “B” is the intensity of GFAP while “b” is the average of these values. For this region, the intensity correlation quotient (ICQ) value is 0.205 (P_sign test_<0.001), indicating partial immunocolocalization (**M**). Scale bars: 50 μm for all.

**Figure 3.**
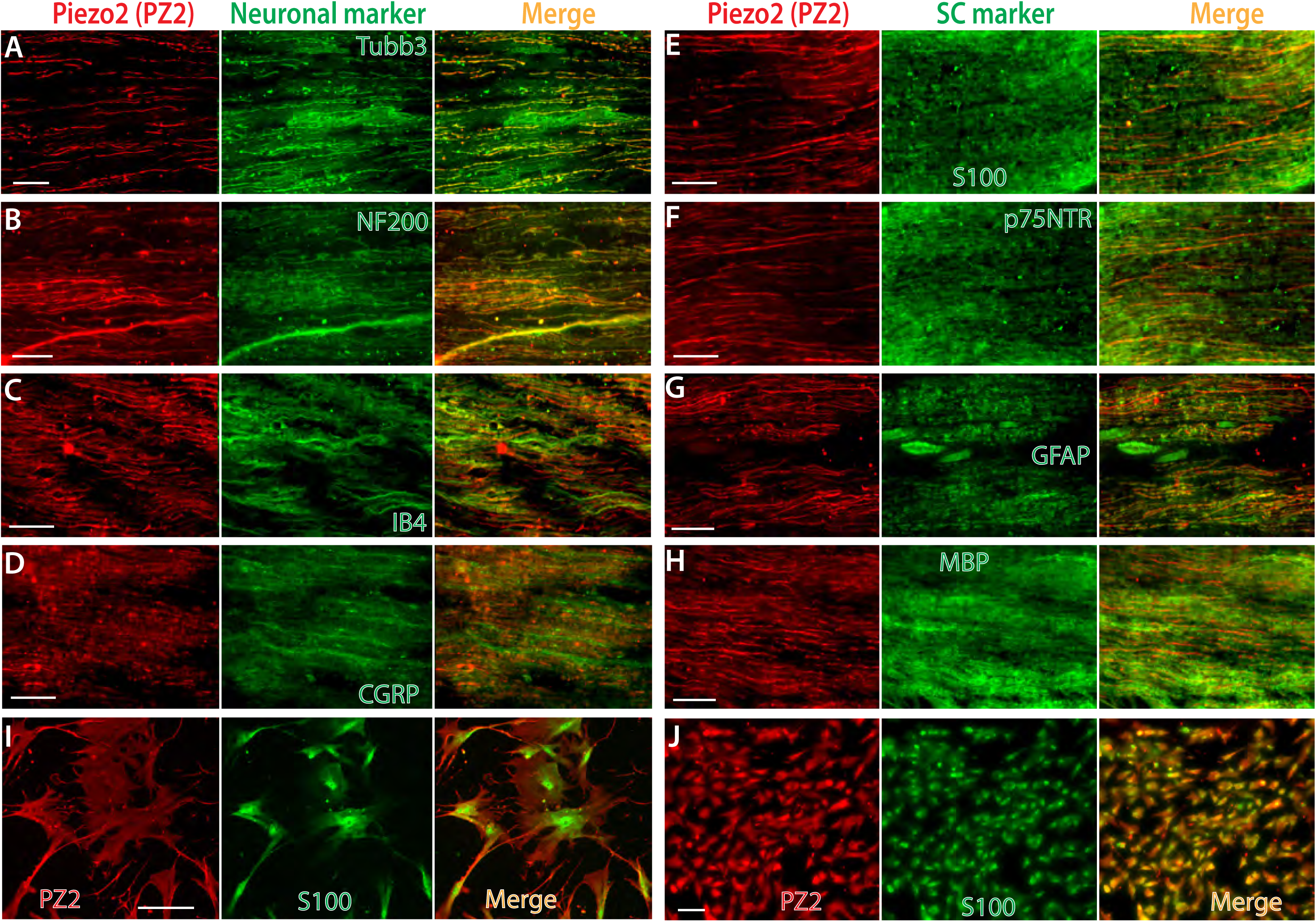
IHC delineation of Piezo2 (PZ2) axonal component and Schwann cell expression. Representative montage images on sciatic nerve cryosections show double immunostaining (IS) of Piezo2 (red) with a selection of neuronal markers (green), including Tubb3 (**A**), NF200 (**B**), IB4 (**C**), and CGRP (**D**). Representative montage images on sciatic nerve cryosections show double-IS of Piezo2 (red) with glia markers (green), including S100 (**E**), p75NTR (**F**), GFAP (**G**), and MBP (**H**). Representative montage images on human Schwann cells (**I**) and isolated Schwann cells (**J**) from rat sciatic nerve show double-IS of Piezo2 (red) with S100 (green). Scale bars: 50 μm for all.

Next, we performed IHC using various established glial cell markers for the satellite glial cells (SGCs) and Schwann cells (SCs), including glutamine synthetase (GS), glial fibrillary acidic protein (GFAP), S100, and myelin basic protein (MBP) [58]. This was performed in order to rule out Piezo2 expression in the perineuronal glial cells, which produce ring-like immunopositivity that resembles NKA1*α* staining patterns. Unexpectedly, IHC staining revealed Piezo2-IR in the S100-, GFAP-, and GS-positive perisomatic glial population, indicating Piezo2 expression in SGCs and/or SCs (**Fig. 2H-L1**) since both express those glial markers [54]. This could be expected on the basis that SGCs may represent a population of developmentally arrested Schwann cells [23]. To further verify the Piezo2 expression in the perineuronal glia, ICA was performed to analyze the immunocolocalization between Piezo2 and GFAP. Overlaid images of Piezo2 with GFAP showed colocalizations of two immunopositivity (**Fig. 2L1**), and the ICA plots of Piezo2 and GFAP also clustered along both positive and negative axes with ICQ 0.06-0.26 (*p* < 0.01∼0.001, n=10), indicating Piezo2 partial colocalization in perineuronal glia cells (**M**). No apparent co-labeling was observed between Piezo2 and MBP which is a major constituent of periaxonal myelin laminae and a myelinating Schwann cells (mySCs) marker. This suggested that Piezo2 was expressed by non-myelinating SCs (nmSCs) but not mySCs (**Fig. 2J**). A search of deposited microarray data in the GenBank GEO Profile corroborated our finding since Piezo2 transcripts, as well as Piezo1 transcripts, have been detected in the cultured human Schwann cells (https://www.ncbi.nlm.nih.gov/geoprofiles/?term=Piezo2+schwann) and SGCs in a recent publication [27]. This provides support for our finding that Schwann cells express Piezo2.

#### 3.2.2. Detection of Piezo2-IR in sciatic nerve, spinal cord, and skin

We next characterized Piezo2 expression in the DRG-PSNs innervating tissues. Results showed that high Piezo2 immunostaining signals were detected in the sciatic nerve, which co-stained with neuronal markers of Tubb3, NF200, IB4, and CGRP (**Fig. 3A-D**), indicating that Piezo2 was actively transported along the afferent axons, including myelinated fibers and C/A*δ* fibers. Notably, sciatic nerve Piezo2 immunostaining displayed Schwann cell components since Piezo2 signals were colocalized to Schwann cell markers of S100, p75NTR, GFAP, and, to a less extent, MBP (**Fig. 3E-H**). ICC colabeling of Piezo2 with S100 on human SCs and isolated SCs from rat sciatic nerve verified Piezo2 expression in SCs (**Fig. 3I, J**).

The spinal cord DH is a key site for integrating and transmitting somatosensory information [2, 31]. GenBank GEO Profile shows both Piezo2 and Piezo1 transcripts detection in spinal cord by microarray (https://www.ncbi.nlm.nih.gov/geoprofiles/?term=Piezo2+spinal+cord) [82], the HUMAN PROTEIN ATLAS shows consensus datasets of Piezo2 and Piezo1 transcripts detection in spinal cord (https://www.proteinatlas.org/ENSG00000154864-PIEZO2/tissue), but no report has found Piezo channel protein expression in the spinal cord [44]. Since IHC revealed actively axonal transporting of Piezo2 to peripheral afferents (**Fig. 3**), we examined whether Piezo2 was also transported to the central terminals innervating the DH. Indeed, Piezo2-IR signals were present in the central terminals of peptidergic and nonpeptidergic neuropil that synapse in the DH of the spinal cord. These are likely transported along axons from DRG-PSNs since DH Piezo2-IR was highly overlaid to presynaptic markers of IB4, CGRP, synaptic vesicle protein synaptophysin (Syp), and synaptoporin (Synpr) (**Fig. 4C-F)** [79]. Additionally, we found extensive Piezo2-IR in the intrinsic spinal cord neurons of both the DH and the ventral horn (VH). Colabeling of Piezo2 with NeuN verified that Piezo2 was expressed in the spinal cord postsynaptic neurons, including DH interneurons and VH motor neurons (**Fig. 4A, B**). DH Piezo2-IR did not present in GFAP-positive astrocytes and Iba1-positive microglia (**Fig. 4G, H**). Finally, Piezo2 was also extensively detected in the brain neurons (**Suppl. Fig. 2**), a finding in agreement with a recent report that shows Piezo2 expression in mouse brain neurons using an independent Piezo2 antibody raised by a Piezo2 C-terminal peptide [69].

**Figure 4.**
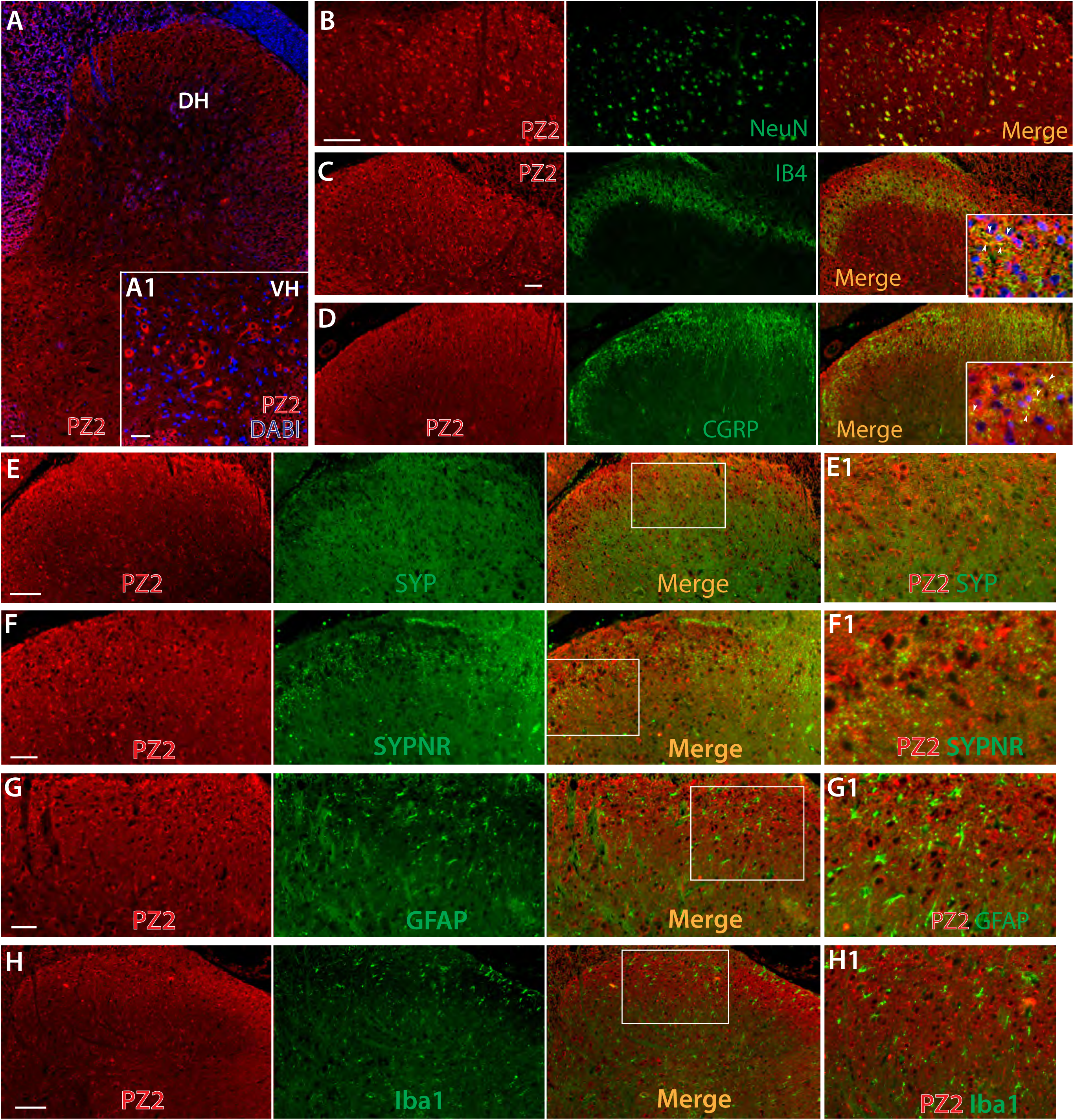
IHC delineation of Piezo2 (PZ2) expression in spinal cord. Representative IHC image shows detection of Piezo2-IR (red) in the spinal cord neurons at all laminae (**A**), with grey matter pseudocolored in blue and a inset showing magnified image, counterstained with Hoechst (blue), of spinal ventral horn (**A1**). Representative montage images of double-IS on the DH regions reveal Piezo2-IR (red) with NeuN (green) (**B**) and a selection of dorsal horn presynaptic markers (green), including IB4 (**C**), CGRP (**D**), Syp (**E**), and Synpr (**F**), showing immunocolocalization (yellow) in the marge images. Insets in the merged images of panels **C** and **D** are higher magnifications of colabeling of Piezo2 (red) with IB4 or CGRP (green) with nuclear counterstained by Hoechst (blue); arrowheads pointing to colocalization (yellow). Representative montage images show negative Piezo2 in GFAP-positive astrocytes (**G**) and Iba1-positive microglia (**H**) of spinal cord sections. The regions within the squares in panels **E**-**H** are shown at high magnification (**E1**-**H1**). Scale bars: 50 μm for all.

Skin Merkel cells express Piezo2, functioning for Merkel-cell mechanotransduction [19, 22, 73]. We also found Piezo2-IR in the Merkel cells of the hindpaw epidermis, which colabeled with CK14, a marker for the basal keratinocytes and Merkel cells (**Fig. 5A, A1, B)** [38, 73]. Data also displayed Piezo2-IR in Meissner’s corpuscles [22] and the afferent terminal nerve bundles in dermis (**Fig. 5B-H)**. Piezo2-IR in the epidermal basal layer keratinocytes was colabeled with S100, which is a well-validated melanocyte (or pigment cells) marker but is negative for Merkel cells (**Fig. 5I, I1)** [46]. Co-labeling of Piezo2 with S100 by ICC on the cultured human epidermal melanocytes verified piezo2 expression in the melanocytes (**Fig. 5K-N**). This indicates that Piezo2 is not restricted to the Merkel cells and epidermal melanocytes also express Piezo2. Additionally, S100-IR was overlaid to the Piezo2-IR signals in Meissner’s corpuscles and Schwann cells surrounding cutaneous afferent nerve bundles within dermis, suggesting that Piezo2 was likely expressed by cutaneous Schwann cells (**Fig. 5H, I, I1, J**). By GFP staining of skin in *Piezo2-GFP-IRES-Cre* knock-in reporter mouse line, Piezo2-GFP fusion was highly visualized in mouse hair skin lanceolate endings [73]. Whether our Piezo2 antibody detects skin lanceolate endings in rat will be investigated in future study.

**Figure 5.**
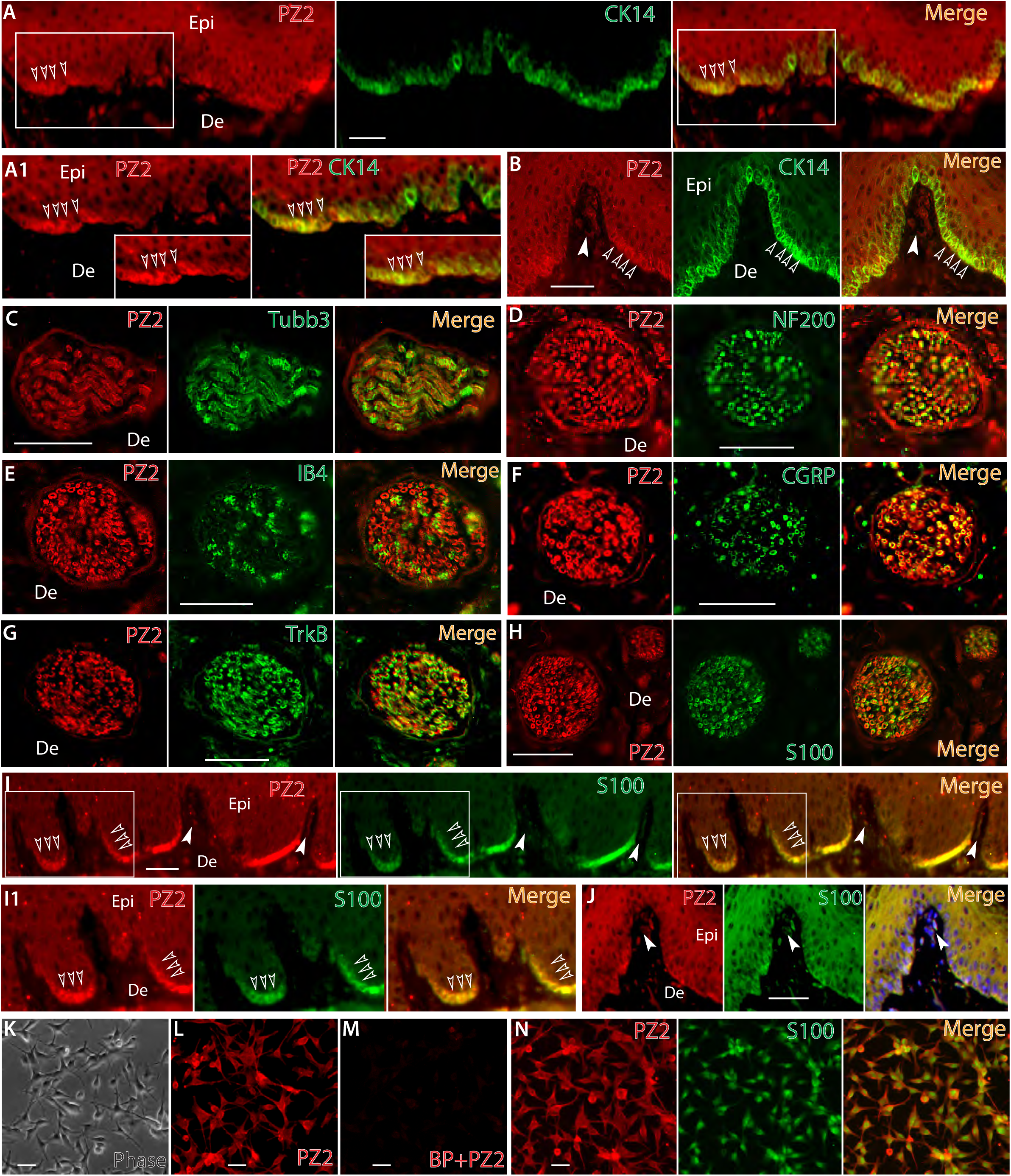
IHC delineation of Piezo2 (PZ2) expression in skin. Representative montage images of hindpaw glabrous skin sections display double immunostaining (double-IS) of Piezo2 (red) with CK14 (green), a maker of epidermal basal layer keratinocytes and Merkel cells, showing colabeling (yellow, empty arrowheads) in the merged image (**A**) with the squared regions shown at high magnification (**A1**). Representative montage images reveal Piezo2-IR in Meissner’s corpuscles (white arrowhead) and immunocolocalization with CK14-positive Merkel cells (empty arrowheads) (**B**). Representative montage images show immunocolocalization of Piezo2-IR with Tubb3 (**C**), NF200 (**D**), IB4 (**E**), CGRP (**F**), TrkB (**G**), and S100 (**H**) in the nerve bundles within dermis. Representative montage images show immunocolocalization of Piezo2-IR with S100 in epidermal basal layer cells (**I**, empty arrowheads. White arrowheads point to Meissner’s corpuscles), with the squared regions shown at high magnification (**I1**). Representative montage images show immunocolocalization of Piezo2-IR with S100 in Meissner’s corpuscles (white arrowheads) (**J**). Human primary epidermal melanocytes (**K**) show Piezo2-IR (**L**) which is completely eliminated by blocking peptide preabsorption before ICC (**M**) and colabeled with S100 (**N**). Scale bars: 25 μm for all.

#### 3.2.3. Confirmation of Piezo2 expression by RT-PCR. RT-PCR

was performed on the total RNAs extracted from 50B11 cells (rat DRG neuronal cells), rat DRG tissue, primary cultured SGCs and Schwann cells prepared from naïve rats, and primary cultured human melanocytes. Results showed amplification of Piezo2 (and Piezo1) transcripts by use of two different pairs of primers specific for Piezo2 (and Piezo1) in these samples, indicating that Piezo2 was expressed in DRG (PSNs), SGCs, Schwann cells, and human melanocytes (**Fig. 6**).

**Figure 6.**
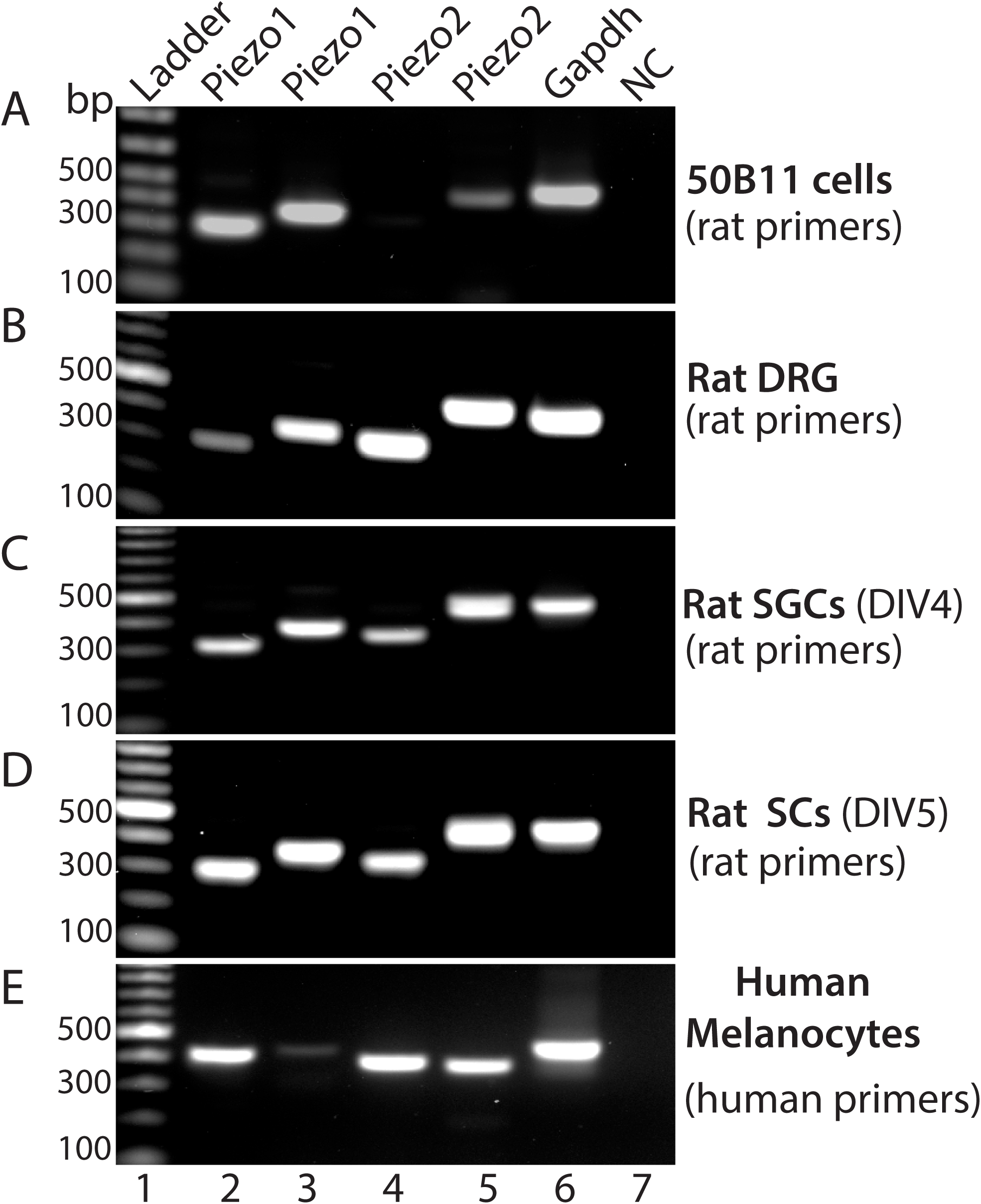
Validation of Piezos (2 and 1) expression by RT-PCR. RT-PCR shows amplification of Piezo1 and Piezo2 transcripts in rat 50B11 cells (**A**), rat DRG (**B**), rat primary cultured SGCs at day of in vitro 4 (DIV, **C**), and rat primary cultured Schwann cells at DIV5 (**D**) by two different primer pairs specific for amplification of rat Piezo1 and Piezo2 transcripts, as well as detection of Piezo1 and Piezo2 transcripts by two different primer pairs specific for human Piezo1 and 2 in human primary cultured melanocytes (**E**). Lane 1: ladder, lane 2-3: Piezo1 amplified by two piezo1 primer pairs (**A-D**, rat; **E**, human), lane 3-4: Piezo2 amplified by two piezo2 primer pairs (**A-D**, rat; **E**, human), lane 7: Gapdh, and lane 8: negative control.

#### 3.2.4. Confirmation with a second Piezo2 antibody

IHC using the alternate Piezo2 polyclonal antibody from Alomone showed a similar profile of Piezo2 immunopositivity in DRG- and TG-PSNs and perineuronal glia, sciatic nerve Schwann cells, Meissner’s corpuscles, and epidermal Merkel cells and melanocytes (**Suppl. Fig. 3**). The observed similarity of patterns in both immunoblots and IHC results between two independent Piezo2 antibodies support the validity of our experimental findings.

### 3.3. Elevated Piezo2 expression in CFA-induced inflammatory pain

Previous studies in human and mouse indicate that Piezo2 is an essential mediator of touch under inflammatory conditions, and inflammatory signals enhance piezo2-mediated mechanosensitivity [18, 62]. We next attempted to determine whether inflammatory pain was associated with alteration of Piezo2 protein expression in DRG and spinal cord. We generated CFA inflammatory pain, confirmed by reduced the threshold for withdrawal from mild mechanical stimulation (von Frey) and hyperalgesia evident with noxious (Pin) mechanical stimulation (**Fig. 7A**) when compared to baseline or CFA rats receiving only saline injection. L4/L5 DRG and lumbar spinal cords were harvested at the 10-day after CFA for western blots (DRG) and IHC (spinal cord). To evaluate the protein expression level of Piezo2 located in the DRG and intracellular trafficking alteration, we separately examined Piezo2 protein levels in the NKA1*α*-enriched membrane fractions versus the NKA1*α*-deficient cytosolic fractions. Multiple bands were noted upon immunoblotting and canonical Piezo2 (∼310KDa) and ∼250-KDa bands, which were clearly separated, were significantly increased in both membrane and cytosols from the DRG ipsilateral to CFA injection, compared to controls (**Fig. 7B, C**).

**Figure 7.**
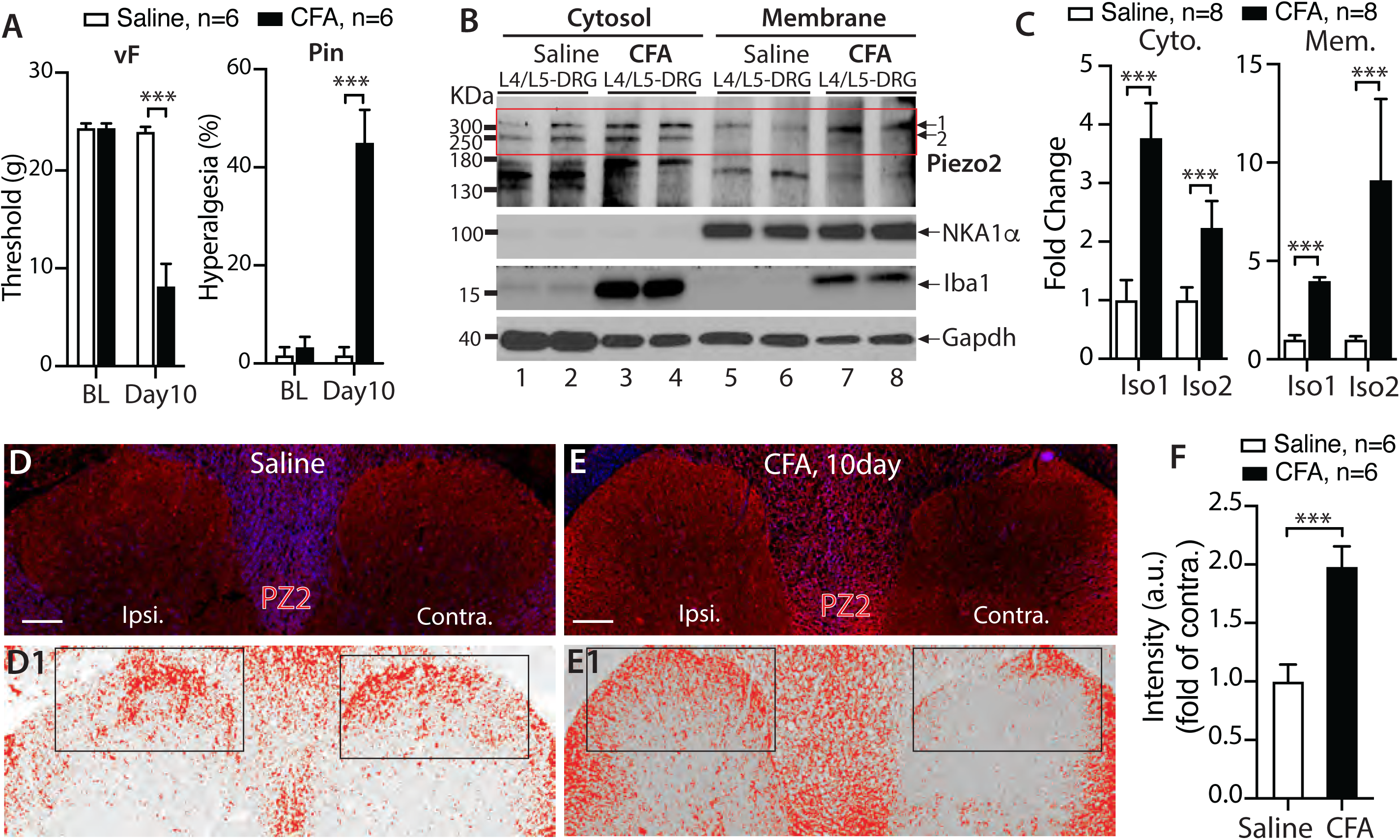
Aberrant Piezo2 (PZ2) expression in CFA-induced inflammatory pain. CFA rats developed mechanical allodynia (vF) and hyperalgesia (Pin) **A**); *** denotes *p*<0.001, unpaired, two-tailed Student’s *t*-test for vF and Mann–Whitney test for Pin, compared between groups after CFA. NKA1*α*-deficient cytosolic fractions and NKA1*α*-enriched membrane fractions were extracted from the DRG (pooled L4/L5) at 10 days after CFA or saline injection, and subjected to immunoblotting (IB) as shown in the representative IBs of Piezo2, Iba1, NKA1*α*, and Gapdh of cytosol (**B**, left) and membrane fractions (**B**, right), respectively. The densitometry of canonical piezo2 (∼310KDa, iso1) and putative iso2 (∼250KDa) outlined by a rectangle were analyzed and summarized in bar charts (**C**); \*\*\**p*<0.001, unpaired, two-tailed Student’s *t*-test. Piezo2 intensities in DH of saline (**D**) and CFA (**E**) were inverted; the upper and lower threshold optical intensity of Piezo2-IR adjusted to encompass and match the IR that appears in red (**D1** and **E1**) with the rectangles positioned over laminae territory throughout the mediolateral axis on the contralateral (contra.) and ipsilateral (ipsi.) sides, and quantified as described in Method. Scale bars: 100 μm for all. The integrated density (product of area and density) calculated by use of ImageJ, and fold change (ratio of ipsi./contra.) summarized in the bar charts (**F**). *** denotes *p*<0.001 by two-tailed unpaired Student’s *t*-test.

Since the spinal cord DH receives innervation from DRG-PSNs, we next evaluated by IHC whether CFA pain was associated with altered Piezo2 expression in the DH. For quantitative comparison, we normalized Piezo2-IR intensity on the side ipsilateral to CFA by using the contralateral side in the same section as the control [14, 57], assuring that all IHC preparation and image capture parameters are identical in the same cross-section. To validate the use of general IR intensity of the DH as a quantitative indicator of protein expression in sensory neurons, we compared IB4 and CGRP staining in the DH from SNI rats, in which quantitatively decreased IB4 and CGRP immunoreactive intensity in sensory neurons ipsilateral to the SNI injury is expected [37, 80]. As shown in **Suppl. Fig. 4**, control rats showed symmetrical patterns of IB4 and CGRP staining in the superficial DH neuropil, whereas 4-week after SNI, quantitative evaluation revealed ipsilateral IB4 and CGRP IR intensity down to 50±4% and 15±3% of contralateral controls, respectively. We next performed Piezo2 IHC on the sections prepared from saline- and CFA-injected rats, and quantitative analysis of the Piezo2-IR intensity ratio between ipsilateral to contralateral DH revealed an enhanced Piezo2-IR in the ipsilateral side compared to the contralateral side in CFA animals (**Fig. 7D-F**).

### 3.4. Increased DH Piezo2-IR intensity in multiple pain conditions

To provide insight into whether DH Piezo2 is also elevated in other pain models, we examined previously archived FFPE axial sections of the lumbar spinal cords from control rats and from rats subjected to neuropathic pain models of SNL, SNI, and TNI (4wk post injury) [37, 57-59, 80, 81], as well as MIA-OA pain (5wk after MIA knee injection), at which time the animals had developed typical hypersensitivity to mechanical stimuli (**Fig. 8A-D**). Results showed that DH Piezo2, determined as the ratio of intensity of IR on the ipsilateral side divided by the contralateral side, was significantly increased in SNL, SNI, and TNI, as well as MIA (**Fig. 8E-M)**. These data suggest that DH Piezo2 protein levels are increased in a wide variety of pain models.

**Figure 8.**
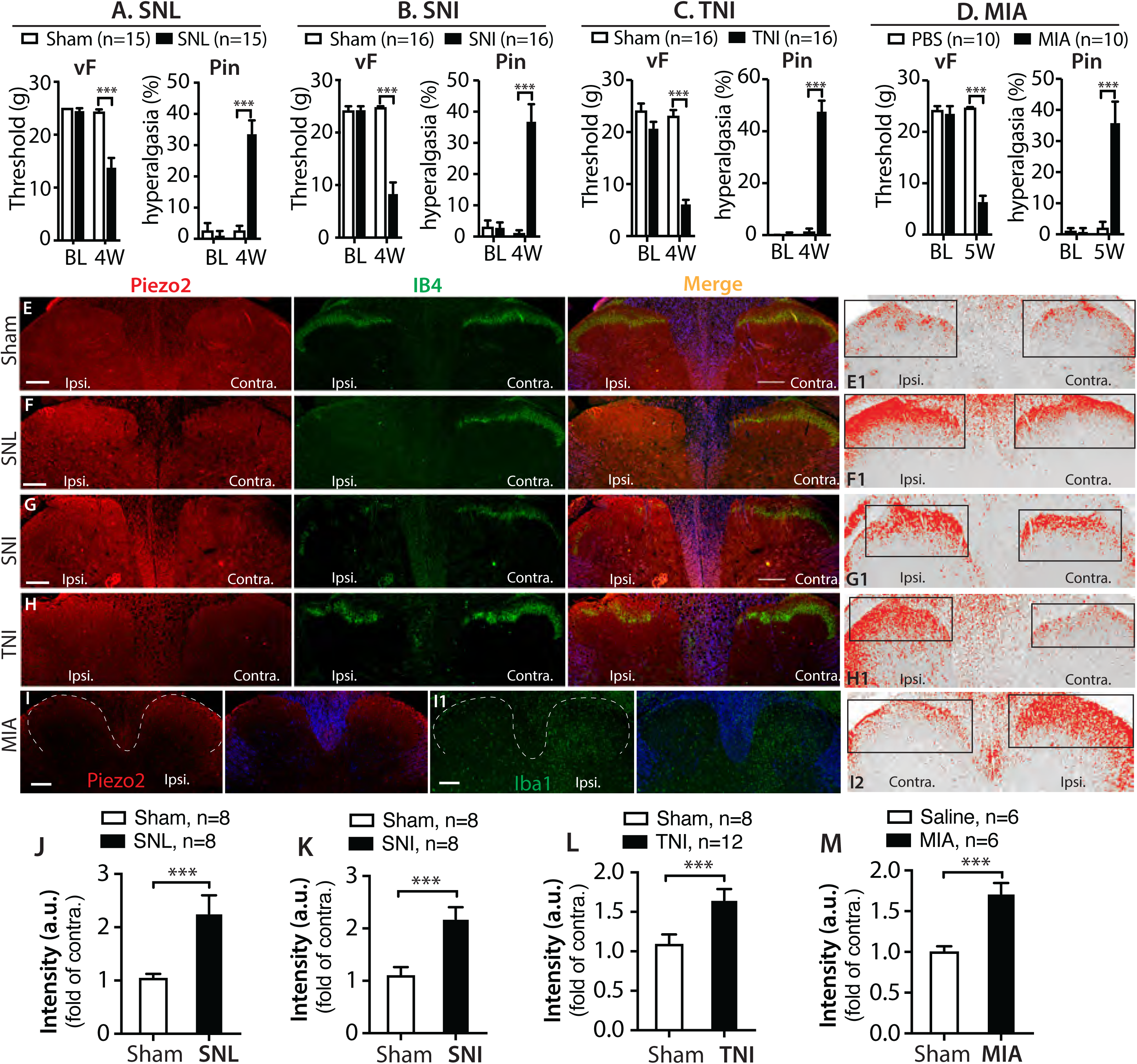
IHC characterization of Piezo2 (PZ2) expression in spinal DH in variety of rat pain models. Bar charts summarize mechanical allodynia (vF) and hyperalgesia (Pin) that encompass various pain models in our previously published studies, as indicated (**A**-**D**). Representative IHC montage images on the DH show double immunostaining (IS) of Piezo2 (red) with IB4 (green) in control (**E**), SNL (**F**), SNI (**G**), and TNI (**H**) rats, and DH Piezo2-IR intensities from the presentative IHC images are inverted, and optical threshold adjusted encompass and match the IR that appears in red (**E1**-**H1**), respectively; and quantified as described in Method. The rectangles positioned over laminae territory throughout the mediolateral axis on the contralateral (contra.) and ipsilateral (ipsi.) DHs. The grey matters in the merged images are pseudocolored in blue. Representative IHC images of Piezo2 (red) (**I**) and Iba1 (**I1**) from two adjacent sections from MIA-OA rat display apparent increased Piezo2-IR in parallel with microgliosis at the ipsilateral side to MIA; white dash-lines outline the dorsal horns (left panels), and grey matters pseudocolored in blue (right panels). Scale bars: 200μm for all. Quantitative comparison of DH Piezo2-IR intensity between ipsi. and contra. sides are analyzed by the Image J (see Methods and suppl. Fig. 5), and the integrated intensity (product of area and density) calculated, and fold change (ratio of ipsi./contra.) presented as the bar charts in SNL (**J**), SNI (**K**), TNI (**L**), and MIA (**M**). *** denotes p<0.001 by two-tailed unpaired Student’s *t*-test.

## 4. Discussion

We present data here showing that Piezo2 is extensively expressed in the sensory neurons and non-neuronal cells in rat peripheral sensory pathways. The major findings include: 1) Piezo2 is expressed by majority of DRG-PSNs, consistent with the recent reports by ISH or IHC showing Piezo2 expression in majority or all DRG-PSNs of mice [69, 83]; 2) Location of Piezo2 expression extends from peripheral terminals in the skin to central presynaptic terminals in the spinal DH; 3) At least four putative Piezo2 isoforms are likely present in DRG, although further biological validation remains to be established; 4) Piezo2 is additionally expressed by DH postsynaptic neurons, motor neurons, and brain neurons; 5) Piezo2 is detected in peripheral non-neuronal cells, including perineuronal glia which are composed of SGCs and nmSCs, sciatic nerve and cutaneous SCs, and skin epidermal melanocytes, as well as Merkel cells; and 6) Spinal DH Piezo2-IR intensity, an indicator of protein levels, was significantly increased in multiple pain conditions, including CFA inflammatory pain, neuropathic pain, and MIA-OA pain. These observations indicate that Piezo2 function related to sensation may involve intimate and sophisticated interactions of PNS neuron-glia and axon-SCs, as well as presynaptic and postsynaptic sensory circuits in the spinal cord.

Piezo2 has emerged as a target of considerable interest in pain research and has mostly been studied in PSNs and Merkel cell biology, touch sensation, and innoxious mechanical pain [6]. However, a consensus is not yet established regarding its role in pain pathogenesis. For example, Piezo2 is persuasively described as a LTMR touch receptor expressed in large PSNs, but new studies indicate that all DRG-PSNs express Piezo2, including IB4 non-peptidergic nociceptors which contain the population of neurons specifically mediating mechanical pain response and CGRP neuron which have been proposed to mainly mediate noxious heat response [61]. Mice PSN-selective Pizeo2 deletion develop tactile allodynia, while other study reports that PSN-selective Piezo2 knockout in mice impairs touch but sensitizes mechanical sensitivity [83]. Piezo2 may also integrate mechanical and thermal cues in vertebrate mechanoreceptors [84]. Function of Piezo2 in pain has been interpreted based on the early experimental data showing that PSNs express only Piezo2 but absent in Piezo1; however, recent studies show that functional Piezo1 is also expressed in PSNs [40, 51, 70]. The Piezo isoforms are not known to form Piezo1-Piezo2 hybrid, but co-expression of Piezo1 and Piezo2 in sensory neurons raises an interesting issue of potential functional interactions between these channels. The apparent similarity in functional properties suggests the possibility of Piezo1 and Piezo2 to cooperate via synergistic or negative interaction [33, 84] or provide redundancy for each other in mechanotransduction and pain pathogenesis.

Detection of Piezo2 protein expression in the spinal cord has not been described previously [44], although credent Piezo2 transcript detection in spinal cord has been reported [82]. Our new data show that Piezo2 expression in PSNs extends from peripheral terminals in the skin to central presynaptic terminals in the DH. Additionally, Piezo2 is abundantly expressed by somata of spinal cord neurons, including superficial and deeper DH neurons and ventral horn motor neurons. Somatosensory information is transmitted from primary afferent fibers in the periphery into the central nervous system via the DH that functions as an intermediary processing center for this information. In general, innocuous touch transmitted by A*β*-LTMR afferents projects to the deep DH LTMR recipient zone (LTMR-RZ) and synapse with interneurons in the deeper DH laminae, while noxious afferents and high threshold mechanoreceptors (HTMRs) terminate predominantly in laminae I–II [2]. Therefore, Piezo2 may play unexplored roles in normal and possibly pain sensory processing of spinal cord. Importantly, we found that the DH Piezo2-IR intensity, an indicator of protein levels, was significantly increased in multiple pain conditions, including neuropathic pain, CFA inflammatory pain, and MIA-OA pain. This may imply the redistributed and upregulated PSN-Piezo2 in the DH central terminals, as have been reported that some pronociceptive channels, such as Na_V_1.7, Na_V_1.8 and Na_V_1.9 or TRPV1, are redistributed at the nerve endings during neuropathic or inflammatory conditions [28, 32, 36, 65]. Future investigation will determine whether increased DH Piezo2 is from redistributed in the central afferent endings or from upregulation of Piezo2 in the DH postsynaptic neurons ipsilateral to peripheral damage.

Peripheral non-neuronal cells play a key role in the induction and maintenance of persistent mechanical pain [41]. Here, we show that Piezo2 is expressed extensively in the PNS non-neuronal cells of various types. The skin acts as a complex sensory organ [41, 60, 85], and epidermal keratinocytes along with Merkel cells, Langerhans cells, and melanocytes express sensor proteins that regulate the neurocutaneous system and participate in nociception and mechanotransduction [41]. Non-neuronal skin Merkel cells express Piezo2, which is required for Merkel-cell mechanotransduction [19, 22, 73]. We detected Piezo2-IR in the skin epidermal Merkel cells and provide evidence that the epidermal melanocytes also express Piezo2 [41]. Skin melanocytes are dendritic cells and derived from skin Schwann cell precursors that origin from neural crest cells (NCC). Melanocytes form tight contacts with cutaneous nerves, respond to mechanical stretch [3, 13, 25, 30], and appear around nerve fascicles after damage [50]. Pressure-sensing Merkel cells are thought of NCC origin [64], but new observations show that Merkel cells originate from epidermal progenitors but not NCCs [66]. Moreover, we found that PNS glia, including SGCs and SCs, express Piezo2. Piezo2-IR was colocalized to S100-positive dermal afferent nerve bundles and terminal fibers, suggesting that Piezo2 is expressed in the cutaneous SCs. Sensory neurons and SGCs/SCs, the two principal cell types of the PNS, interact intimately and are critical for normal PNS functions and pain pathogenesis during nerve injury and inflammation. Peripheral SCs can sense and transduce mechanical signals involving mechanosensation [8, 52], and the specialized cutaneous nociceptive nmSCs in skin determine the sensitivity threshold for mechanosensation [1]. Therefore, peripheral nerve SCs-Piezo2 may play hitherto unexplored roles in sensory processing and mechanotransduction; for instance, by regulating signaling within SCs initiated by mechanical stimulation [8, 11, 52].

We also detected extensive Piezo2 expression in rat brain neurons, consistent with a recent independent report showing Piezo2 expression in adult mouse brain neurons [69]. It is not wholly surprising to identify Piezo2 in brain since the *Piezo* gene is originally discovered in brain and dubbed as *Mib (*Membrane protein induced by Aβ) [55]; and study using brain neuronal analogue N2A cells discovers *Mib* as a mechanosensor and subsequently renamed as Piezo2 [15].

**5**.In conclusion, Piezo2 is widely expressed by neuronal and non-neuronal cells of the PNS and by neurons in the spinal cord and brain. Further study is needed to verify Piezo2 functions in these various cells. Overall, our results suggest that Piezo2 is not only involved in detection of external mechanical stimuli in the skin, but may also serve to detect internal mechanical cues in the nervous system or link to other unexplored intercellular and intracellular signaling pathways in the pain circuits [9]. Our findings provide additional sites for further investigation to fully understand the roles of Piezo2 in sensation and related to pain pathogenesis.

## Acknowledgments

This research was supported by a grant from the Department of Veterans Affairs Rehabilitation Research and Development I01RX001940 (to QH), a National Institutes of Health grant R61NS116203 (to HY and QH), a National Institutes of Health grants NS108278 (to CLS), and an MCW NRC grant FP00016291 (to HY).

