## Supplemental_Figures for "Piezo2 mechanosensitive ion channel is located to sensory neurons and non-neuronal cells in rat peripheral sensory pathway: implications in pain"

  

**Suppl**. **Fig**. **1**. **ICC
of Piezo2 (PZ2) in DRG dissociated culture.**

Representative montage images of Piezo2
immunostaining by ICC on DRG dissociated culture at Days In Vitro (DIV) =1 show immunoreactivity of Piezo2
(red) in neuronal somata and their axons (white arrowheads), as well as smaller
glia-like cells and their neurites (empty arrowheads) (**A**).  
  Representative montage images of
ICC double immunostaining on DRG dissociated culture (DIV=0.25) show
Piezo2 (red, white arrowheads) immunopositivity in all-sized neurons and co-stained
with S100-positive (green, empty arrowheads) glia cells (**B**). Representative montage images of ICC double immunostaining on DRG dissociated culture (DIV=2) show immunocolocalization
of Piezo2 (red) with GFAP-positive (green) glia cells (**C**). Scale
bars: 100 mm for all.

  

**Suppl**. **Fig**. **2**. **IHC
detection of Piezo2 expression in brain neurons.**

Representative
IHC images show detection of Piezo2
(PZ2) expression (red) in the
neurons of prefrontal cerebral cortex (Cx1) (**A**), lateral cerebral cortex
(Cx2) (**B**), and hippocampus (Hip) (**C**, **D**) areas of brain FFPE
sections from naïve adult rat. Representative montage images (vertical) show absence
of PZ2 (red) in the GFAP-positive astrocytes (green) in naïve rat brain (**E**). Scale bars: 25 μm for all.

  

**Suppl**. **Fig**. **3**. **Characterization
of a different Piezo2 antibody in detection of Piezo2 (PZ2) expression.**

In
DRG lysate, Piezo2 antibody (Alomone)  detects several
bands at approximate 310,
240, 180, and 130 KDa and marked as putative “isoforms” 1-4 of the target
protein, since they are eliminated by preincubation with excess immunogenic peptide (**A**). Representative montage images of double immunostaining
(double-IS) on DRG sections reveal Piezo2-IR (red) and a selection of neuronal
markers (green), including Tubb3 (**B**), IB4 (**C**), CGRP (**D**),
and NKA1a (**E**), showing
immunocolocalization (yellow) in merged images. Representative montage images of double-IS on trigeminal
ganglia (TG) section reveal Piezo2-IR (red) and Tubb3 (green) (**F**). The panels in the right-side of **B**-**D** are the percentage of Piezo2-IR neurons
overlaid to Tubb3-positive neurons (**B**1), as well as IB4- (**C1**) and CGRP-positive neurons (**D1**) overlaid to Piezo2-IR neurons and the
numbers are the counted Piezo2-IR neurons (red) and marker-IR neurons (green)
in double labeling sections. Representative montage images of double-IS on
DRG sections display Piezo2-IR (red) and a selection of glial cell markers
(green), including GFAP (**G**) with
the region within the square shown at high magnification (**G1**), S100 (**H**), MBP (**I**), showing
immunocolocalization (yellow) in merged images. Representative montage images of double-IS on sciatic nerve sections reveal Piezo2-IR (red) and S100 (**J**)
and GFAP (**K**), showing immunocolocalization (yellow) in merged images. Representative montage images of double-IS on the hindpaw glabrous skin display Piezo2-IR
(red), NF200 (green) (**L)** and IB4 (green) (**M**), CK14 (green) (**N**,
epidermal Merkel cells pointed by empty arrowheads and Meissner's
corpuscles pointed by white arrowheads), S100 (**O**,
epidermal melanocytes pointed by empty arrowheads and Meissner's
corpuscles pointed by white arrowheads), showing
immunocolocalization (yellow) in merged images. Scale bars: 50 μm for all.

  

**Suppl**. **Fig**. **4**. **Validation
of the methods to quantify DH immunostaining intensity**.

IB4
and CGRP immunolabeled fluorescent intensities in spinal DH of sham control (**A,** top) and SNI (**B,** top) were inverted; the upper and lower threshold
optical intensities of IB4 and CGRP signals adjusted to encompass and match the
IR that appears in red (**A, B,** bottom), respectively; and quantified as
described in Method. The rectangles (**A, B,** bottom) positioned over
laminae territory throughout the mediolateral axis on the contralateral and ipsilateral
DHs. Scale bars: 100 mm for
all. The integrated density
(product of area and density) calculated by use of ImageJ, and fold change (ratio
of ipsilateral/contralateral) summarized in the bar charts (**C**). \* and \*\*\*
denotes *p*<0.05 and *p*<0.001 by two-tailed unpaired Student’s *t*-test. c, contralateral and i, ipsilateral.
